# The effect of hypoxia on *Daphnia magna* performance and its associated microbial and bacterioplankton community: a scope for Genotype x Microbial community interactions upon environmental stress ?

**DOI:** 10.1101/2023.02.09.527849

**Authors:** Manon Coone, Isabel Vanoverberghe, Shira Houwenhuyse, Chris Verslype, Ellen Decaestecker

## Abstract

The depletion of oxygen as a result of increased stratification and decreased oxygen solubility is one of the most significant chemical changes occurring in aquatic ecosystems as a result of global environmental change. Hence, more aquatic organisms will be exposed to hypoxic conditions over time. Deciphering the effects of hypoxia on strong ecological interactors in this ecosystem’s food web is critical for predicting how aquatic communities can respond to such an environmental disturbance. Here, (sub-)lethal effects of hypoxia and whether these are genotype specific in *Daphnia*, a keystone species of freshwater ecosystems, are studied. This is especially relevant upon studying genetic responses with respect to phenotypic switches (G x E interactions) upon environmental stress. Further, we investigated the effect of hypoxia on the *Daphnia* microbial community to test if the microbiome plays a role in the phenotypic switch and tolerance to hypoxia. For this, two *Daphnia* genotypes were exposed for two weeks to either hypoxia or normoxia and host performance was monitored together with changes in the host associated and free-living microbial community after this period. We found G x E interactions for some of the tested *Daphnia* performance traits. The microbial community responded to hypoxia stress with responses in the bacterioplankton and in the *Daphnia* associated microbial community with respect to species richness and community composition and structure. The latter response was different for the two genotypes suggesting that the microbiome plays an important role in G x E interactions with respect to hypoxia tolerance in *Daphnia*, but further testing (e.g. through microbiome transplants) is needed to confirm this.

## 1 Introduction

The depletion of oxygen is one of the most significant chemical changes currently occurring in freshwater ecosystems as a result of global environmental change. While hypoxia is common as a seasonal disturbance, the duration, spatial scale and frequency have increased in the past few decades and are expected to increase as a result of climate change (Diaz and Rosenberg, 2008, Gilbert et al., 2005, Paerl et al., 2011, Goto et al., 2012). This is concerning as hypoxia has a strong effect on these freshwater ecosystems. The main drivers of deoxygenation are rising water temperature, stratification and anthropogenically induced eutrophication. Warmer surface water holds less soluble oxygen, which in its turn results in increased thermal stratification due to the increasing density difference with deeper colder water depths. This increasing separation reduces circulation between different depths of the water column. Reduced oxygen exchange between the atmosphere and the water causes a further decrease in oxygen amount. Excess nutrient runoff, primarily nitrogen and phosphorus, that enters the water column often leads to plankton blooms, which upon death sink to the bottom and increased oxygen consuming decomposition activity leads to hypoxic conditions in the deeper water depths. This reduction in dissolved oxygen promotes nutrient release from bottom sediments into the surface water (North et al., 2014), enabling the production of phytoplankton blooms. As hypoxic waters and phytoplankton blooms act as reinforcing factors of each other, their co-occurrence is often reported and reinforces that nutrient excess is one of the major causes of increasing oxygen depletion (Zhang et al., 2011). The effect of increasing deoxygenation is 2.75 to 9.3 times larger in freshwater lakes than in oceans, with losses in dissolved oxygen (D.O.) concentration of 5.5% in upper regions and 18.6% in lower regions of 393 studied freshwater bodies between 1980 and 2017 (Jane et al., 2021). Water is considered hypoxic when the dissolved oxygen levels are less than 2 mg/L, since these levels are linked with harmful effects on fish and zooplankton (Vanderploeg et al., 2009).

Deciphering the effects of hypoxia on strong ecological interactors in the food web of freshwater ecosystems, such as the zooplankter *Daphnia magna*, is thus essential to predict if and how aquatic communities can respond to such a disturbance. When dissolved oxygen becomes depleted in water ecosystems below organismal physiological tolerances, it can (in)directly impact aquatic communities and natural resources. *Daphnia* is adapted to hypoxia via a series of physiological (Pirow et al., 2001) and biochemical (Gerke et al., 2011) adaptations, each with their own advantages and costs (Larsson and Lampert, 2011, Galic et al., 2019). Mobile organisms, like *Daphnia*, are less prone to direct lethal effects compared to sessile organism (Díaz and Rosenberg, 1995), but they risk indirect effects of hypoxia when fleeing to more oxygenated regions. On the one hand, hypoxia-induced growth reduction (Seidl et al., 2005, Kobayashi, 1982, Eby and Crowder, 2002) is beneficial for diffusive oxygen transport pathways (Pirow et al., 2004, Pirow and Buchen, 2004, Seidl et al., 2005), but on the other hand, it is detrimental since the smaller body size implies a diffusive bypass in the haemolymph circulation (Pirow et al., 2004, Pirow and Buchen, 2004) which reduces the effectiveness of the circulatory system’s ability to resist oxygen overload in body tissues in regions with high D.O. concentrations (Seidl et al., 2005). Depending on the *Daphnia* species, hypoxia can impact reproduction impairments (Seidl et al., 2005, Lyu et al., 2013a). Over a wide range of atmospheric oxygen concentrations, *Daphnia* can control their oxygen metabolism and metabolic phenotype (Weider and Lampert, 1985, Lee et al., 2022). For example, to avoid fish predation, *Daphnia* uses vertical migration to seek refuge into hypoxic regions with D.O. concentrations lethal for fish (Larsson and Lampert, 2011, Hanazato et al., 1985, Decaestecker et al., 2002). During hypoxia exposure, *Daphnia* upregulate hemoglobin synthesis resulting in a higher hypoxia tolerance (Pirow et al., 2001) and a red phenotype (Seidl et al., 2005, Gorr et al., 2004, Zeis et al., 2013), making *Daphnia* lose their transparent appearance and visual advantage for predation (Larsson and Lampert, 2011). The amount of dissolved oxygen has a significant impact on the population composition as not all genotypes are as effective at surviving in hypoxic conditions, which leads to selection and shifts in the genotype composition of the *Daphnia* population (Hutchinson, 1957). While respiration rate is genotype independent (Weider and Lampert, 1985), hemoglobin synthesis is genotype dependent (Weider and Lampert, 1985). Genotypes with a lower hypoxia tolerance have an enlarged vulnerability to predation causing changes in community structure, variations in the distribution of species and reduction in biodiversity. During periods of hypoxia, low hypoxia tolerant genotypes may be forced to move to oxygen regions, which are still tolerant for fish (Weider and Lampert, 1985), but the hypoxia-induced red phenotype makes them more visible and prone to predation. Further research is needed into the physiological or ecological mechanisms underlying a *Daphnia* population tolerance to hypoxia. Such as the mediation of hypoxia tolerance through the microbiome.

We here hypothesize that microbial communities change upon hypoxia and may indirectly affect host metabolism or phenotypic effects upon selective uptake or rejection of particular microbiota by the host. Microbial data in combination with host performance effects are needed for a comprehensive understanding of hypoxia induced effects on hosts and their microbiomes. Because of their high phenotypic plasticity in response to environmental stressors, short generation time, small size, large number of eggs per clutch, easy manipulation, absence of ethical concerns in experiments and a fully sequenced genome, *Daphnia* is a model organism for eco-(toxico)logical research (Ebert, 2005, Miner et al., 2012, Ebert, 2022). Moreover, recent results show that *Daphnia* has a flexible microbiome increasing its tolerance upon environmental stress (Macke et al., 2017b, Akbar et al., 2022). They have an easy-to-manipulate gut microbiome and it is possible to cause strong disturbances in the microbial community upon sterilization (Callens et al., 2016). The *Daphnia* microbial composition is dynamic and differs between gut and outer body parts (Sison-Mangus et al., 2015, Callens et al., 2018, Qi et al., 2009). In contrast to the whole microbial community of *Daphnia*, the gut microbial community is dominated by only a small number of bacterial groups (e.g. Comamonadaceae, Flavobacteriacea, Burkholderiaceae, *Aeromonas, Limnohabitans, Pedobacter, Ideonella* and *Pseudomonas* (Akbar et al., 2020, Akbar et al., 2022, Macke et al., 2017a, Cooper and Cressler, 2020, Qi et al., 2009). The environment and host genotype, which influence the microbial community composition and functionality to achieve a balanced state, can cause these interactions to be temporary and susceptible to selection (Macke et al., 2020, Akbar et al., 2022). It has been shown that depriving *Daphnia* of its microbiota is detrimental to its fitness and the association between microbial imbalance and disease states is becoming clear, also in *Daphnia* (Rajarajan et al., 2022, Bulteel et al., 2021). *Daphnia* gut microbial community responses have been detected upon exposure to environmental stress (Houwenhuyse et al., 2021), for example upon exposure to antibiotics (Callens et al., 2018, Akbar et al., 2020) and parasites (Bulteel et al., 2021, Rajarajan et al., 2022). Another specific example of the effects of a microbiome-related reaction to environmental conditions in *D. magna* was described for toxic cyanobacteria. Eutrophication, rise in temperature and depleted oxygen concentrations, due to global warming, increases toxic cyanobacterial blooms (cyanoHABS), which is associated with reduced fitness of *D. magna* and a changed gut microbial community (Macke et al., 2017a). A gut microbial community transplant experiment showed that cyanobacteria sensitive *D. magna* genotypes became more tolerant after being transplanted with the gut microbiota from a tolerant *D. magna* genotype (Macke et al., 2017a).

We here investigated whether the *Daphnia* associated microbial community can mediate responses to hypoxia with a particular focus on genotype x microbial community x environmental (G x M x E) interactions, as these may mediate microbiome mediated evolutionary responses for *D. magna* populations during times of hypoxia in the environment. Therefore, we first investigated the (sub)lethal effects of hypoxia in *Daphnia* and whether these are genotype specific. Next to the study of the *Daphnia* G x E interaction in response to hypoxia stress, we investigated if the microbiome is relevant in phenotypic responses towards hypoxia. Two *Daphnia* genotypes were exposed for two weeks to either hypoxia or normoxia and survival, growth and reproduction were monitored together with changes in host associated microbial communities. The first hypothesis that was tested is that the *D. magna* genotype mediates tolerance towards hypoxia exposure (G x E). The second hypothesis tested is that particular microbial communities are associated with hypoxic conditions (M x E) and are potentially relevant for *Daphnia* tolerance to hypoxia. To determine the effect of hypoxia on the microbial community of *Daphnia*, the microbial composition of the gut and body of two genotypes was characterized via amplicon sequencing. In addition, bacterioplankton samples of the medium were taken to investigate whether free-living microbial communities released by the *Daphnia* host also reflected a shift with decreasing oxygen levels and if responses were different than the host associated microbial communities (body and gut). We here investigated whether host genotype x microbial community interactions occur upon hypoxia to further test microbiome-mediated genotype specific effects in hypoxia tolerance in *Daphnia*.

### 2 Materials and methods

### 2.1 *D. magna* and *Chlorella vulgaris* cultivation

Throughout this study, two *D. magna* genotypes were used: the KNO 15.04 and the F genotype. Genotype KNO15.04 originates from a small (350m^2^), fishless, mesotrophic pond in Knokke, at the Belgian coast (51°20′05.62″ N, 03°20′53.63″ E). The F clone is a standard genotype used in ecotoxicological tests, obtained from the Barrata lab in Barcelona and originally isolated in Scotland (Barata et al., 2017). All *D. magna* stock genotypes were cultured and maintained for many generations in the Aquatic Biology lab (IRF life sciences lab, KU Leuven department Kortrijk, Belgium). For the experiment, three maternal lines of these stock lines were set up per genotype to exclude maternal effects between individuals of the same genotype. The maternal lines were established by collecting and continuing every second (or third) brood during two or more generations. *D. magna* used in the experiments were obtained from eggs of the second or third brood, given that these are better quality than first brood offspring. Cultures were kept at a density of one *D. magna* individual per 50 mL in filtered tap water (Greenline e1902 filter) at a constant room temperature of 19±1°C and under a 16:8h light-dark cycle. They were fed three times a week with 200.10^3^ cells/ml of the unicellular green algal species *Chlorella vulgaris. Chlorella vulgaris* cultures were cultured under sterile conditions in 2L jars with Wright’s Cryptophyte medium (Guillard and Lorenzen, 1972) at a constant room temperature of 20±2°C and under a light-dark cycle of 16:8h. To prevent bacterial contamination, a 0.22µm filter was present at the in- and output of the aeration system to supply CO2 and to remove oxygen. Magnetic stirrers were used to ensure a constant mixing in order to avoid precipitation of the algae. Fluorescence-activated cell sorting was used to measure the cell density of the algal cultures (using FACS Verse, Biosciences).

### 2.2 Experimental set-up

To unravel if different *D. magna* genotypes show a different sensitivity towards hypoxia exposure, the two *D. magna* genotypes (KNO 15.04 and F) were exposed to a hypoxic and normoxic exposure. Female *D. magna* carrying parthenogenetic eggs in their brood pouch, at a stage of ≥ 48h after egg laying, were isolated for three maternal lines per genotype. Once the *D. magna* juveniles were released from the brood pouch, they were individually transferred to a 50 mL Falcon tube containing filtered tap water. Per maternal line, ten *D. magna* juveniles for each of the three maternal lines used per genotype were exposed to either a hypoxic or a normoxic exposure for fourteen days. The hypoxic exposure was achieved using Biospherix C-chambers with a ProOx controllers, where a 2% air oxygen level was reached in the chambers using regulated inflow of nitrogen gas under pressure (1.7 mbar). Dissolved oxygen content (mg/L), oxygen saturation (%), temperature (°C) and pressure (hPa) were measured daily to check the stability of the hypoxic exposure using the Hach HQ40d multi-meter and optical dissolved oxygen sensor. The dissolved oxygen in the medium decreased gradually, reaching a stable level of ± 1.83 mg/L after two days. The experimental exposures were kept constant throughout the experiment with a temperature of 19 ±1°C and a 16-8h light-dark cycle. To account for differing light incidence and potential differences between the hypoxia chambers, falcons were randomized daily. After the transfer to the normoxic or hypoxic exposure, survival and fecundity (day of first and second brood, and brood amount) were monitored daily. Body size of 5 *D. magna* individuals per combination of maternal line and exposure was measured at days 3, 7, 10 and 14 of the exposures using a BMS microscope camera and BMS PIX software. The length of a *D. magna* was measured from the top of the eye to the base of the apical spine. Starting from day two of the experiment, 200.10^3^ cells/mL of autoclaved *Chlorella vulgaris* were administered every other day.

### 2.3 Amplicon sequencing

At the end of the experiment, surviving individuals were dissected and *D. magna* guts and bodies were collected separately to determine the microbial community composition of each genotype and body part via amplicon sequencing. In addition, bacterioplankton samples of the medium were taken by filtering 200 mL of the medium over a 0.22 µm filter. Samples were collected on ice (to prevent microbial community shifts and to preserve DNA) in an Eppendorf tube containing 10 µL sterile Milli-Q. For each exposure, individuals were pooled per genotype and maternal line. Amplicon sequencing was performed according to Houwenhuyse et al., 2021. DNA was extracted by the Qiagen PowerSoil DNA isolation kit and dissolved in 20 µL MilliQ water. To determine the total DNA yield, 1 µL of sample was used in an Invitrogen Qubit dsDNA HS assay. Increased specificity and amplicon yield were obtained by using nested PCR. The entire 16S rRNA gene was amplified for 30 cycles (98 °C for 10 s; 50 °C for 45 s; 72 °C for 30 s) with the Life Technologies SuperFi high fidelity polymerase and the EUB8F and 1492R primers on 10 ng of template. The PCR product was purified with the QIAquick PCR purification kit. In a second amplification round, 5 µL (20-50 ng) of the PCR product was amplified for 30 cycles (98 °C for 10 s; 50 °C for 5 s; 72 °C for 30 s) with the 515F and 806R primers to obtain amplicons of the V4-region with a dual index. The latter two primers contained an 8-nucleotide barcode, as well as an Illumina adapter at their 5’-end. PCRs were performed in triplicate and were pooled and gel-purified for each sample with the QIAquick gel extraction kit. To prepare an equimolar library, an Applied Biosystems SequalPrep Normalization Plate was used to normalize amplicon concentrations, after which the library was pooled. Amplicon sequencing was performed using a v2 PE500 kit with custom primers on the Illumina Miseq platform, which resulted in two 250-nucleotide paired-end reads for each of the 36 samples.

### 2.4 Statistical analyses

All data analysis was performed using R 4.0.4. To select the models with the best combination of variables, the Akaike information criterion (AIC) was used. For survival data, A log-rank test was performed using the ‘survdiff’ function (survival package in R) to determine whether there was a significant difference in survival probability between groups. Survival probability over time for different groups was visualized by plotting Kaplan-Meier curves using the ‘ggsurvplot’ function (survminer package in R). For body size, differences between groups were determined by performing an Analysis of Variance (ANOVA) using the ‘Anova’ function (car package in R) on a generalized linear model (GLM) and contrasts between specific groups was analyzed with a Tukey post-hoc test. When taking maternal lines into account as a random factor, a linear mixed-effects model (lmer function, lme4 package in R) was used on normally distributed data or a generalized linear mixed effects model (glmer function, lme4 package in R) was used when the data was not normally distributed. Body size was compared over time and for each time point separately. The best model was determined by having the lowest AIC. Differences in body size between groups was visualized by making boxplots using the ‘ggplot’ function (ggplot2 package in R). Total fecundity was analyzed with a linear mixed-effects model, controlling for an unbalanced design with a restricted maximum likelihood estimation.

We processed DNA sequences in accordance with Callahan et al., 2016b. Sequences were trimmed on both paired ends (the first 10 nucleotides and starting at position 180) and filtered (maximum of 2 expected errors per read). The high-resolution DADA2 method, which relies on a parameterized model of substitution errors to discriminate sequencing errors from actual biological variation, was used to predict sequence variations (Callahan et al., 2016a). After that, chimeras were removed from the data set. Using the SILVA v138 training set, a naïve Bayesian classifier was used to assign taxonomy. ASVs that were classified as “chloroplast” or “cyanobacteria” or that had no taxonomic assignment at the phylum level were eliminated from the data set. Following filtering, a total of 738198 reads—an average of 19951.3 reads per sample—were obtained, with the majority of the samples having more than 9000 reads. ASVs were pooled at the order level, and orders accounting for less than 1% of the readings were disregarded in order to visualize the bacterial orders that varied between the treatments. Measures for alphadiversity of the microbial communities within the different exposures, genotypes and sample types (ASV richness and Shannon Index) were calculated using the vegan package in R (Bellier, 2012). Prior to analyzing alphadiversity, all samples were rarified to a depth of 9500 reads, based on the number of reads per sample. A generalized linear model (GLM), assuming a Poisson distribution of the data, was used to investigate the effects of sample type (gut, body, or bacterioplankton), oxygen exposure (normoxia or hypoxia), genotype (KNO 15.04 or F), and any possible interactions on ASV richness with maternal line as random factor. The ‘emmeans’ function with a ‘Tukey’ adjustment from the emmeans R package was used to perform pairwise comparisons between significant variables and their interactions. Principal Coordinates Analysis with the phyloseq package in R was used to calculate and plot weighted and unweighted Unifrac distance matrices in order to compare variations in microbial community composition and structure (beta diversity) between variables. Using the Adonis2 function in the vegan package in R, the effect of oxygen exposure, genotype, sample type, and all potential interactions on β-diversity were evaluated through a permutation MANOVA. Obtained p-values were adjusted for multiple comparisons through the control of the false discovery rate (FDR). To identify which bacterial classes significantly differed between the exposures and sample types, ASVs were grouped at the class level, and classes representing <1% of the reads were removed. Differential abundance analyses were then performed with the Bioconducter package DESeq2 (Love et al., 2014).

## 3 Results

### 3.1 Effects of hypoxia on *Daphnia* performance traits

The two-way interaction between *Daphnia* genotype and oxygen exposure (comparison normoxia versus hypoxia) was significant for survival (Supplementary Figure 1; Cox proportional hazard model: interaction *Daphnia* genotype x oxygen exposure: X^2^ = 11; df= 3; p = 0.01). When the two genotypes were pooled, the *Daphnia* individuals which were subjected to normoxia had a higher survival rate compared to the ones in hypoxia (Supplementary Figure 2B; Cox proportional hazard analysis: treatment: X^2^=7.1; df=1; p=0.008). Consistent with the significant interaction between genotype and exposure treatment, the tendency was that survival was slightly more reduced in the KNO 15.04 than in the F genotype in hypoxia versus normoxia (Figure 1). A two-way interaction between genotype and oxygen exposure over time could also be seen for body size (p<0.01). The hypoxia exposure had a significant overall negative effect on *Daphnia* growth over time (Figure 2, p<0.001). The general increasing effect of hypoxia on growth over time was mainly attributed to the stronger limiting effect of hypoxia on the growth of the KNO 15.04 genotype (Figure 3B right panel, p<0.0001) compared to the F genotype *Daphnia* individuals (Figure 3A left panel, p=0.5). At day 14, KNO 15.04 *Daphnia* individuals that were exposed to hypoxia were 15.27% smaller compared to *Daphnia* of the same genotype who were exposed to normoxia. For the F genotype, there was only a reduction in size of 0.28% between normoxia and hypoxia at day 14 (Figure 2). Notable, there was no differential growth between the two *Daphnia* genotypes in normoxia (Figure 2: right panel: p=0.28). The F genotype *Daphnia* individuals became larger than the KNO 15.04 genotype *Daphnia* individuals in hypoxia over time (Figure 2 left panel: p<0.01). Both genotypes had a clear red phenotypic appearance in hypoxia (Supplementary Figure 3) and of the *Daphnia* surviving until day 14, the proportion of hypoxia exposed reproducing *Daphnia* (5%) was significantly lower than normoxia exposed *Daphnia* (40%) (Figure 3A; p<0.01). Also, the number of eggs in the first clutch was significantly lower in hypoxia compared to normoxia exposed *Daphnia* (Figure 3B; p<0.05). Genotype did not influence the number of reproducing *Daphnia*, nor the amount of produced first clutch eggs. However, the KNO 15.04 genotype had a higher percentage of *Daphnia* individuals carrying a second clutch compared to the F genotype in normoxia (Figure 3C; p<0.05). Clutch size did not differ between the genotypes. Not a single *Daphnia* individual reproduced a second time in hypoxia (Figure 3 C&D).

**Figure 1.**
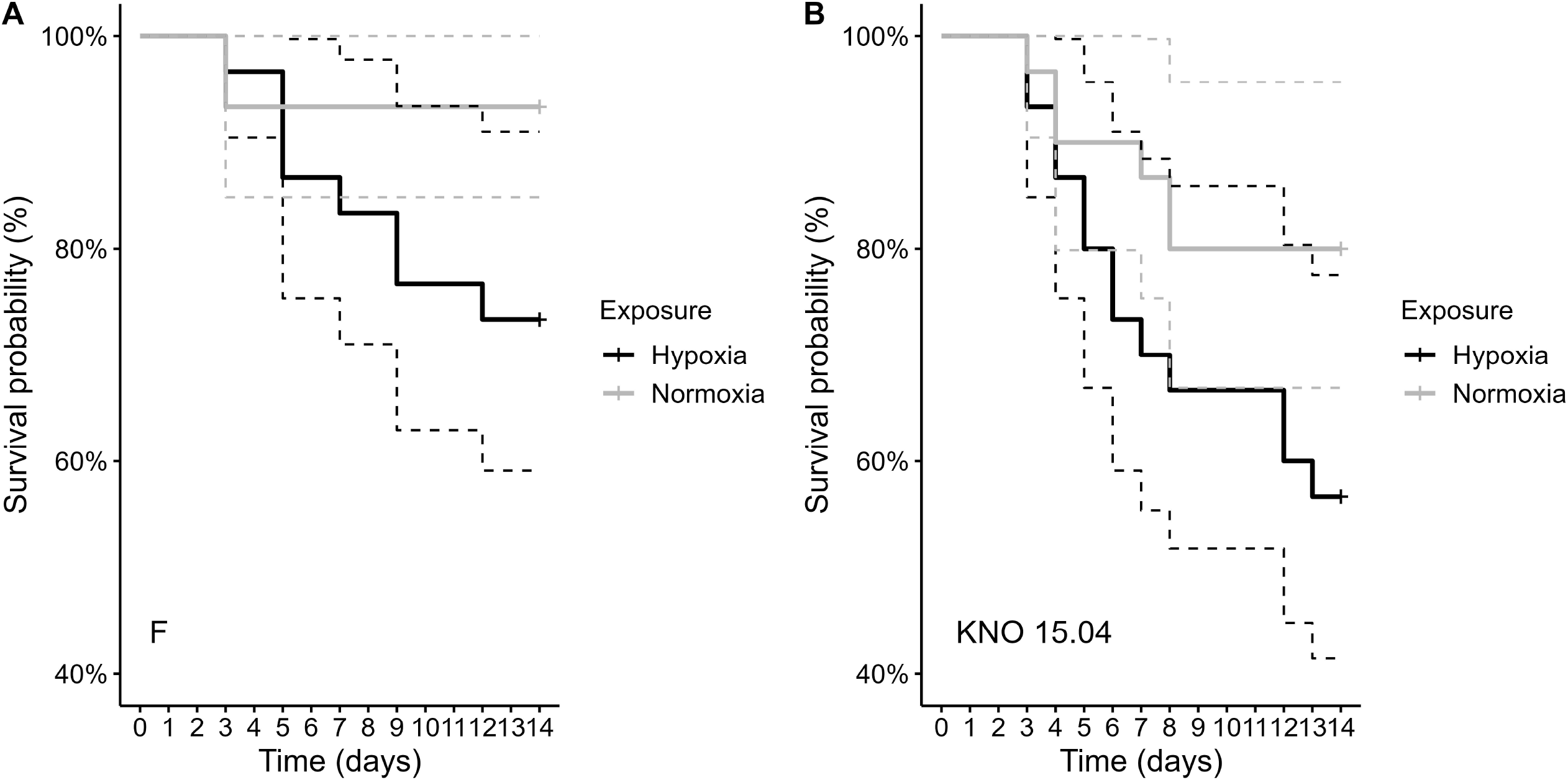
Comparison of survival of the F genotype (A) and KNO 15.04 genotype (B) in the hypoxia (black line) or normoxia (grey line) exposure. Dashed lines represent the 95% confidence intervals. Sample size was n=30 (10 individuals * 3 independent replicates) for each genotype * exposure combination.

**Figure 2.**
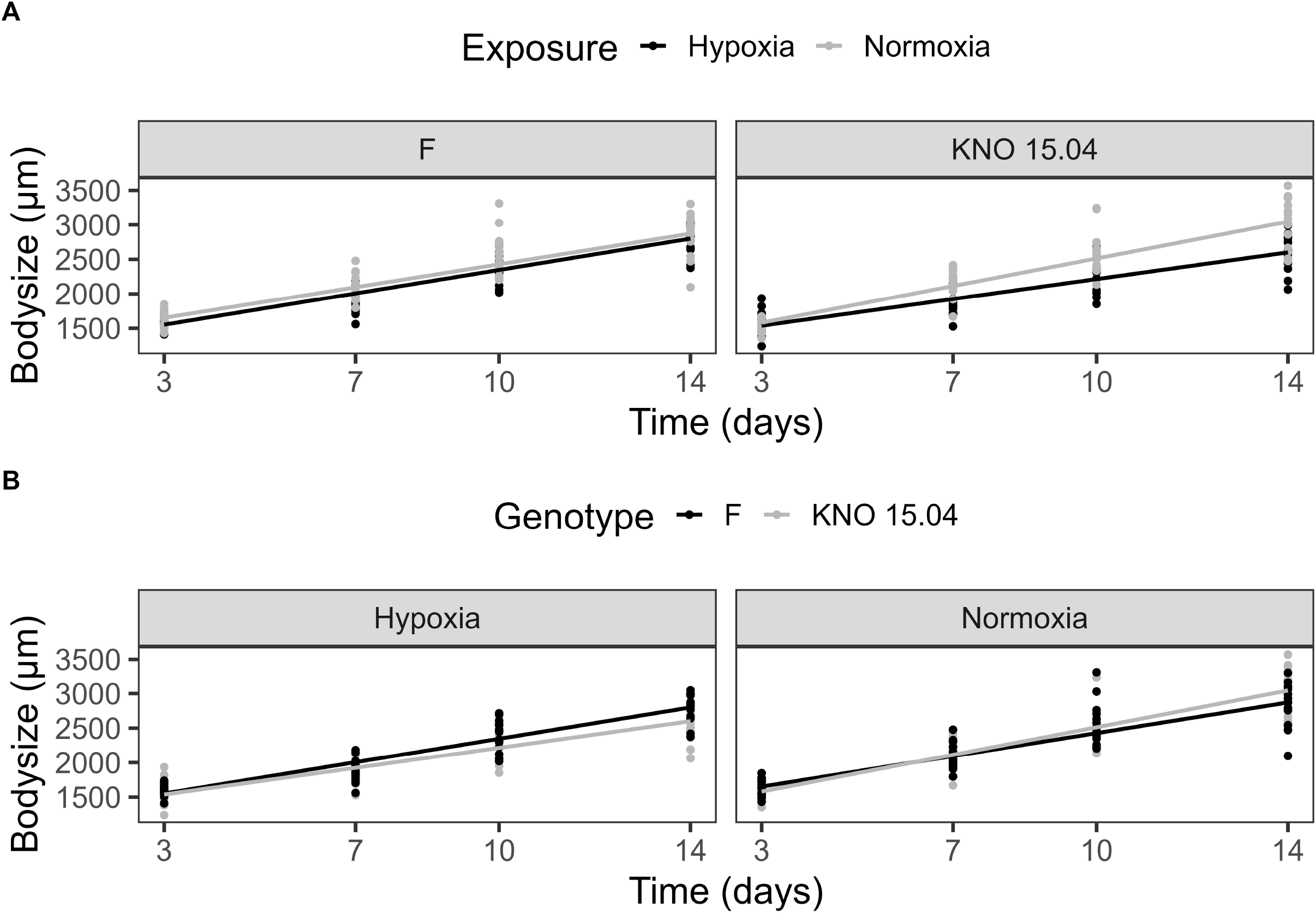
Upper panel: Exposure effect on the body size of the F (left) and KNO 15.04 (right) genotype during hypoxia (black line) or normoxia (grey line) exposure over 14 days. Lower panel: Genotype effect in the hypoxia and normoxia exposure: Black lines correspond to body size of the F genotype and grey lines to body size of the KNO 15.04 genotype. Dots represent individual Daphnia body size data.

**Figure 3.**
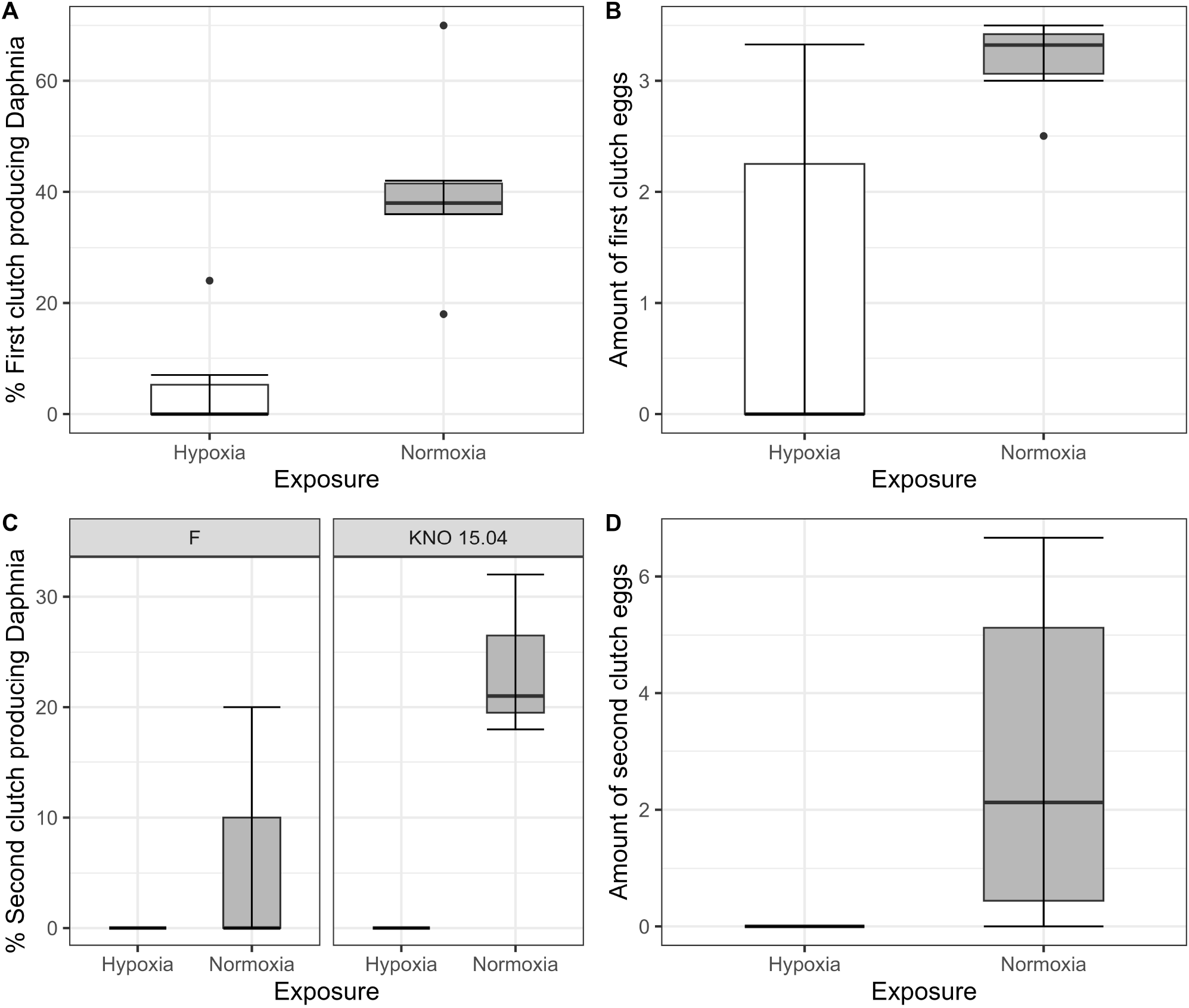
Fecundity of Daphnia genotypes F and KNO 15.04 under hypoxia (white boxplots) and normoxia (grey box plot). A) Percentage of first clutch producing individuals of the Daphnia surviving up to 14 days. B) Size of first clutch in the two exposures. C) Percentage of second clutch producing individuals of the Daphnia surviving up to 14 days of the F and KNO 15.04 genotype, left and right panel, respectively. D) Size of second clutch in the two exposures. The box plot includes 50% of the data from the first to the third quartile, the whiskers extend to the minimum and maximum data within the 1.5 interquartile range and the dots represent single outlier data points outside that range.

### 3.2 Microbial community responses to hypoxia

The species richness (SR) of the complete dataset showed a two-way interaction between exposure and sample type (p<0.05), with a higher SR in bacterioplankton compared to body and gut samples (bacterioplankton-body samples: p<0.0001, bacterioplankton-gut samples: p<0.001, ANOVA) when pooling the two genotypes in hypoxia. This was reflected in a higher amount of Actinobacteria (Wald test: BPK-body: padj<0.0001; BPK-gut: padj<0.05) and a lesser amount of Gammaproteobacteria (Wald test: BPK-body: padj<0.0001; BPK-gut: padj<0.0001) in the bacterioplankton samples compared to the *Daphnia* body and gut in hypoxia (Figure 4). The difference in SR between bacterioplankton and the other sample types was not present in normoxia (p=0.09). However, also in normoxia Gammaproteobacteria were less present in the bacterioplankton compared to the gut samples (Wald test: padj<0.05). In addition, the bacterioplankton contained more Bacteroidiota than the *Daphnia* gut samples (Figure 4, Wald test: padj<0.05) and more Verrucomicrobiae than the *Daphnia* body (Wald test: padj<0.01) and gut samples (Wald test: padj<0.05) in normoxia (Figure 4). When taken relative abundances of species into account, the two-way interaction between sample type and exposure disappeared, as the Shannon Index (SI) of the bacterioplankton samples differed significantly not only from gut (p<0.001) and body (p<0.001) samples in hypoxia, but also from gut samples in normoxia (p<0.05).

**Figure 4.**
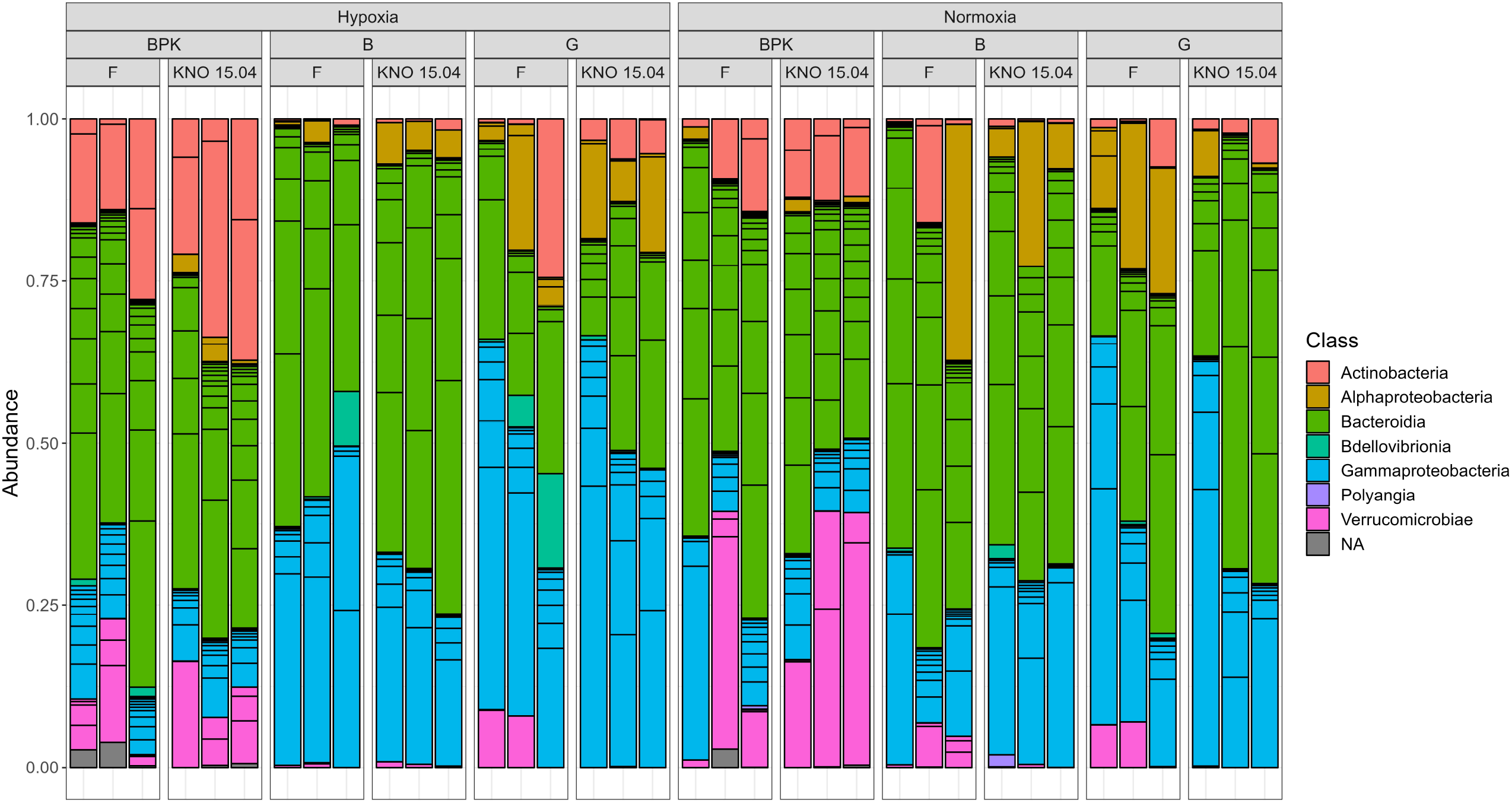
Overview of the relative abundance of different classes of bacteria present in three sample types (Bacterioplankton - BPK, Body – B, Gut – G of the two D. magna genotypes (F versus KNO 15.04).

When looking at the exposure effect on the separate sample types, hypoxia only affected the bacterioplankton community significantly, where there was an enrichment in Actinobacteria (Wald test: padj<0.05) and a reduction in Gammaproteobacteria (Wald test: padj<0.05) and Verrucomicrobiae (Wald test: padj<0.05) (Supplementary Figure 4). No significant exposure effect could be found within the body and gut samples community using the Deseq2 analysis, but when looking at the rare classes (representing less than 1% of the data), it can be seen that the class Polyangia was no longer represented in hypoxia in both the gut and body samples compared to normoxia (Supplementary Table 1). For the gut samples, the classes Acidimicrobiia, Armatimonadia and Kapabacteria were no longer present in hypoxia (Supplementary Table 1). While there were no differences in species richness (p=0.98) nor Shannon Index (p=0.99) between gut and body samples, it should be noted that when investigating rare bacterial classes, the classes Bacilli, Desulfitobacteriia, Acidimicrobiia, Armatimonadia and Kapabacteria were not present in the *Daphnia* body samples, while they were present in the *Daphnia* gut samples (Supplementary Table 1).

The three-way interaction between exposure, sample type and genotype in SI (p<0.05) can be explained by the fact that the two-way interaction between exposure and sample type differed for the different genotypes. This three-way interaction was only borderline significant for SR (p=0.08). In the F genotype there was a two-way interaction between exposure and sample type: sample types did not differ from each other in normoxia, but they did in hypoxia: bacterioplankton samples differed from both body (p<0.05 and p<0.01, for SR and SI, respectively) and gut (p<0.05 and p<0.05, for SR and SI, respectively) samples (Figure 5). In the KNO 15.04 genotype, there was no two-way interaction: the SR of bacterioplankton samples differed from both body and gut samples in normoxia as well as in hypoxia (Normoxia: bacterioplankton-body samples: p<0.0001, bacterioplankton-gut samples: p=0.0001; Hypoxia: bacterioplankton-body samples: p<0.0001, bacterioplankton-gut samples: p<0.001, ANOVA). For SI, the difference between bacterioplankton samples and both body and gut samples in normoxia became smaller in hypoxia to a point were none of the sample types differed significantly from each other when the relative abundances of the species were taken into account (Figure 5: SI: Normoxia: bacterioplankton-body samples: p<0.01, bacterioplankton-gut samples: p<0.05; Hypoxia: bacterioplankton-body samples: p=0.06, bacterioplankton-gut samples: p=0.12, ANOVA). Within the gut samples, there was a significant two-way interaction between exposure and genotype in SR. Within the F genotype, gut samples in hypoxia had a significant lower species richness compared to normoxia (p<0.05). This trend towards a lower species richness and Shannon index upon hypoxia, although not significant, could also be seen in the body samples of the two genotypes, but not in the gut samples of KNO 15.04. An opposite trend could be seen in the bacterioplankton samples, where hypoxia exposure resulted in a higher SR and SI than in normoxia in the F genotype and a higher SR in the KNO 15.04 genotype (Figure 5). In normoxia, the gut of the KNO 15.04 genotype had a significantly lower species richness compared to the F genotype (p<0.05).

**Figure 5.**
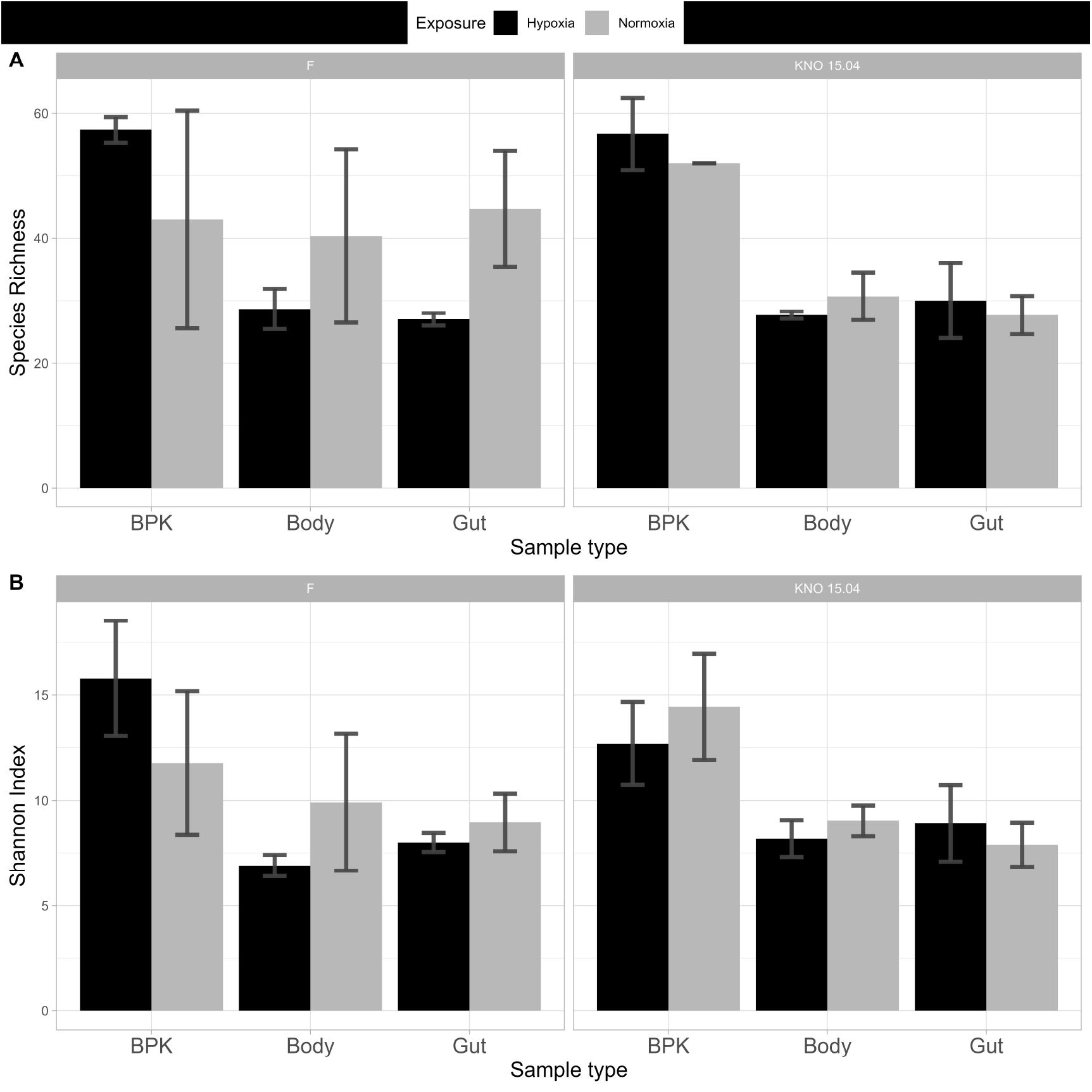
Alpha diversity: (A) Species Richness and (B) Shannon Index of bacterial communities between genotypes and sample types in hypoxia (black) and normoxia (grey). Left panels: F-clone; right panels: KNO 15.04.

For beta-diversity, the composition of the sample types did not differ from one another in normoxia (p=0.19), but hypoxia exposure made the bacterial communities of the different sample types more distinct from each other causing bacterioplankton, body and gut samples to form three distinct clusters, when pooling genotypes (p=0.0001). This distinction between sample types was present in the structure of the bacterial microbiota community in normoxia (p<0.01), but was more pronounced in hypoxia (p=0.0001). The strongest effect could be seen in bacterioplankton samples in which both structure (p<0.05; Supplementary Figure 5A left panel) and composition (p<0.05. Supplementary Figure 5B left panel) differed between normoxia and hypoxia. In the *Daphnia* body samples, the composition of the bacterial community differed between normoxia and hypoxia (p<0.05; Supplementary Figure 5B middle panel), but the overall structure was the same (p=0.35; Supplementary Figure 5A middle panel). There was no difference in composition (p=0.18; Supplementary Figure 5B right panel) or structure (p=0.25; Supplementary Figure 5A right panel) of the gut bacterial communities between hypoxia and normoxia.

When looking at the genotypes separately, the bacterial microbiota of the different sample types did not differ in their overall structure (p=0.29; Figure 6A upper right panel) and composition (p=0.89; Figure 6B upper right panel) in normoxia, but hypoxia significantly impacted both the overall structure (p<0.01; Figure 6A upper left panel) and composition (p<0.01; Figure 6B upper left panel) of the bacterial microbiota causing a more distinct bacterial community per sample type in the F genotype. This change of going from a shared (more common) structure and composition between the sample types in normoxia towards a distinct structure and composition per sample type in hypoxia is less pronounced in the KNO 15.04 genotype in which this is only true for composition (p=0.09 in normoxia and p<0.05 in hypoxia; Figure 6B lower right and left panel, respectively). The overall structure of the sample types of the KNO 15.04 genotype was not significantly impacted by hypoxia as the sample types already significantly differed from each other in normoxia (p<0.01; Figure 6A lower right panel) and remained to differ in hypoxia (p<0.01; Figure 6A lower left panel).

**Figure 6.**
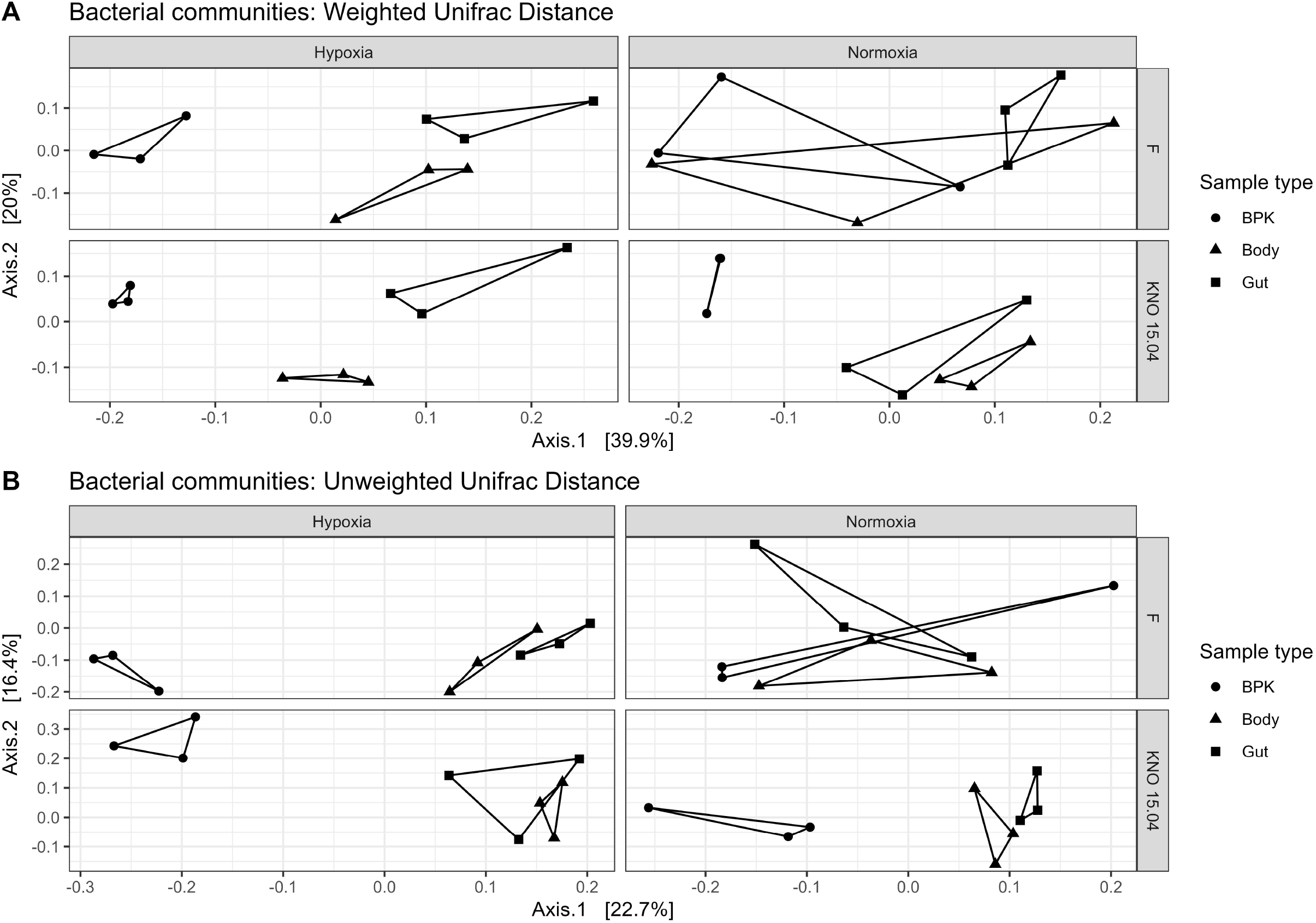
Beta diversity: (A) Weighted and (B) Unweighted Unifrac distance of the bacterial communities for the different genotypes in hypoxia and normoxia. Circles correspond to bacterioplankton (BPK) samples, triangles to body samples and squares to gut samples.

## 4 Discussion

In this study we investigated the effect of a long-term hypoxia exposure on *D. magna* performance and its associated microbial community. The microbial communities studied were these from *Daphnia* body and gut and the free-living bacterioplankton released by *Daphnia*. As hypothesized, hypoxia at an environmentally relevant concentration reduced survival, fecundity and growth of the *D. magna* individuals tested. These results are in line with previous research in *D. magna* (Homer and Waller, 1983) and another *Daphnia* species, *D. similis*, that has intensively been investigated under hypoxia (Lyu et al., 2013a). In comparison with *D. similis*, the *D. magna* genotypes here tested had high survival percentages, which are in line with high survival percentages of *D. magna* in earlier studies (Lyu et al., 2013a, Homer and Waller, 1983). This suggests that *Daphnia* tolerance to hypoxia is interspecific even for *Daphnia* species with a similar body size. Hypoxia here affected mainly reproduction via the amount of first and second clutch producing *D. magna* individuals and the amount of the first clutch eggs produced, which is in line with earlier effect in *D. similis* (Lyu et al., 2013a) and reproduction impairments in *D. magna* exposed to D.O. levels below 2.7 mg/L (Homer and Waller, 1983). But our results contradict (Seidl et al., 2005), where clutch size was unchanged in the first five clutches in *D. magna* acclimated to hypoxia. Hypoxia induced reproduction impairments can affect future generations and the population structure in the aquatic environment. In this study, hypoxia had the largest effect on body size, where a genotype x exposure two-way interaction showed a genotype dependent effect of hypoxia on body size with the KNO 15.04 genotype growing slower compared to the F genotype in hypoxia. Reduction in body size as a result of hypoxia exposure is consistent with other studies in *Daphnia* spp. (Seidl et al., 2005, Lyu et al., 2013a, Hanazato, 1996, Homer and Waller, 1983) and a smaller body size is assumed to improve hypoxia tolerance through better diffusive oxygen-transport processes (Pirow et al., 2004, Pirow and Buchen, 2004). *Daphnia* individuals divert energy from their development, growth and reproduction towards producing hemoglobin to facilitate oxygen uptake under hypoxic stress (Hanazato and Dodson, 1995). The fact that we did not find strong G x E interactions in survival and reproduction may be because we tested two genotypes that showed a relative high hypoxia tolerance. Body size was more affected than survival and fecundity, suggesting that different *D. magna* performance traits are differentially responsive to hypoxia. A similar phenomenon was found for nitrite presence, another environmental stressor linked with water pollution, where reproduction was more affected than survival and molting (Lyu et al., 2013b).

Diet and antibiotics are known traditional environmental factors that shape the gut microbial composition in a way that can be associated with changes in performance traits, also in *D. magna* (Akbar et al., 2020). We here showed that also hypoxia is an environmental factor that affects *Daphnia* associated microbial communities and especially the bacterioplankton that surrounded the *Daphnia* individuals. To determine the effect of hypoxia on the microbial community of *D. magna*, the microbial composition of the gut and body of the two genotypes was investigated. In addition, bacterioplankton samples of the medium were taken to investigate whether microbial communities in the medium experienced a shift with low oxygen levels. Two patterns with respect to selective uptake of microbial strains to obtain tolerance towards a toxic cyanobacterial diet upon the exposure of *Daphnia* to microbial inocula have been proposed by Houwenhuyse et al., 2021: (1) selection of specific beneficial and/or adapted strains, and/or (2) selection for a high strain diversity with complementary gene functions. While support for both these patterns was found for the tolerance of *D. magna* to the toxic cyanobacteria (Houwenhuyse et al., 2021), our results only showed evidence of the first pattern. Important to note is that in our study no extra inocula were added, so the microbial community in the bacterioplankton are the bacterial strains that were removed from the *Daphnia* and were growing in the experimental *Daphnia* medium. In our results, there was a trend towards a reduced alpha diversity in terms of species richness and Shannon index in *D. magna* body tissue and gut upon hypoxia, which can be seen especially in the gut samples of the F genotype where species richness was significantly lower in hypoxia compared to normoxia. This trend was associated with a trend towards an increased alpha diversity in the bacterioplankton samples, causing the bacterioplankton community to differ strongly from gut and body samples in hypoxia while the bacterioplankton community was similar to gut and body bacterial communities in normoxia. Although the F genotype was clearly more responsive to hypoxia than KNO 15.04 with respect to responses in the microbial community, there was no significant G x E interaction on microbial alpha diversity. Studies where *Daphnia* gut and bacterioplankton differed in their microbial community demonstrate the impact of the environmental conditions of *Daphnia* on their interaction with microbial symbionts (Callens et al., 2016, Freese and Schink, 2011). On the one hand, Actinobacteria were found to be more abundant in the bacterioplankton compared to body and gut in hypoxia which reflects a potential expelling effect under hypoxia by *D. magna*. Bacteria of the class Verrucomicrobiae, on the other hand, seem to thrive well under hypoxia as they are more abundant in bacterioplankton in normoxia compared to body and gut samples, while in hypoxia the amount does not significantly differ between sample types. This theory that Verrucomicrobiae are only partially secreted in the medium in hypoxia is supported by the fact that they are less than half as abundant in the bacterioplankton in hypoxia compared to normoxia. The sample type differences in the alpha diversity were translated in the beta diversity. When looking at the difference between communities, hypoxia caused the composition and structure of the bacterial communities from the bacterioplankton, gut and body to change from more similar communities to three distinct community clusters in the F genotype. This could only be seen in the composition of the sample types in the KNO 15.04 genotype, while the differences between the microbial structure of the sample types became larger in hypoxia. The F genotype showed thus a stronger microbial response to hypoxia for both beta- and alpha-diversity.

Our results support the theory that environmental stress can alter host-microbial community relationships and the pattern seems to be host genotype dependent (Akbar et al., 2020). We assume that this effect affects *Daphnia* performance but further testing, e.g. through microbiome transplant experiments, for this is needed. No general microbiome adaptation towards a hypoxia environment was found and although the F genotype showed a stronger response to hypoxia in terms of microbial responses and survival, it was the KNO 15.04 that showed the strongest effects on body size and kept its microbial community equal between the different sample types tested. Also, non-adaptive symbiont loss due to hypoxic stress, similar to the process of symbiont loss in coral bleaching (Johnson et al., 2021), is a hypothesis that we cannot exclude here. Another possibility is that the detected changes in host performance and shifts in microbial communities are due to a change in *Daphnia* metabolism and filtration(feeding) rates (Hanazato and Dodson, 1995, Lee et al., 2022). The metabolic phenotype of *Daphnia* in response to hypoxia was found to be influenced by both micro-evolutionary differences and spatial and temporal environmental heterogeneity of the aquatic environment (Lee et al., 2022, Weider and Lampert, 1985). *Daphnia* subjected to hypoxia as a result of thermal stratification are also exposed to variations in pH, temperature, salinity, conductivity, nutrients and food levels. Studies that consider the interacting impacts of these factors that may also affect metabolism and life history traits are required in order to effectively estimate the effects of lower DO in aquatic environments. It has already been shown that further reductions in life history traits arose in hypoxia, if it is accompanied by food shortage as food shortage and oxygen deficiency cause synergistic effects on the life history of *D. magna* (Hanazato, 1996).

In conclusion, hypoxia reduced survival, body size and reproduction in *D. magna* differentially with the strongest genotype specific effect reflected in *Daphnia* body size. Alongside impairements in *D. magna* performance traits, hypoxia induced the composition and structure of the bacterioplankton, and the *Daphnia* associated microbial communities (gut and body) to shift into distinct communities. Within the gut and body samples a trend towards a lower alpha diversity in hypoxia was found, which was associated with a higher alpha diversity in the surrounding bacterioplankton and this effect was different for the two genotypes. This finding is relevant in the context of host acclimatization and evolutionary potential upon climate change, which is the primary cause of hypoxia and is predicted to worsen over the next decades, inducing effects in the zooplankton and its associated microbial community and in turn affecting the biodiversity of the natural freshwater systems with effects for the quality of drinking water.

## 6 Author Contributions

MC and ED designed the experiment. MC performed the experiment and IV performed extractions and sample processing for sequencing. MC analyzed the data, with help from SH. MC drafted the manuscript and managed revisions with input from ED and CV.

## 8 Acknowledgments

We are grateful for the assistance of Dzhamilyat Kurbanova and Alec De Buyck during the experimental work. We thank Amruta Rajarajan, Luc De Meester, Robby Stoks en Koenraad Muylaert for the stimulating discussions and Eline Beert and Jonas Blockx for the technical help with the hypoxia chambers.

## Data Availability Statement

The generated and analyzed datasets for this study will be made available after acceptance on NCBI.

