## Supplementary figures 1-5 and table1 for "The effect of hypoxia on *Daphnia magna* performance and its associated microbial and bacterioplankton community: a scope for Genotype x Microbial community interactions upon environmental stress ?"

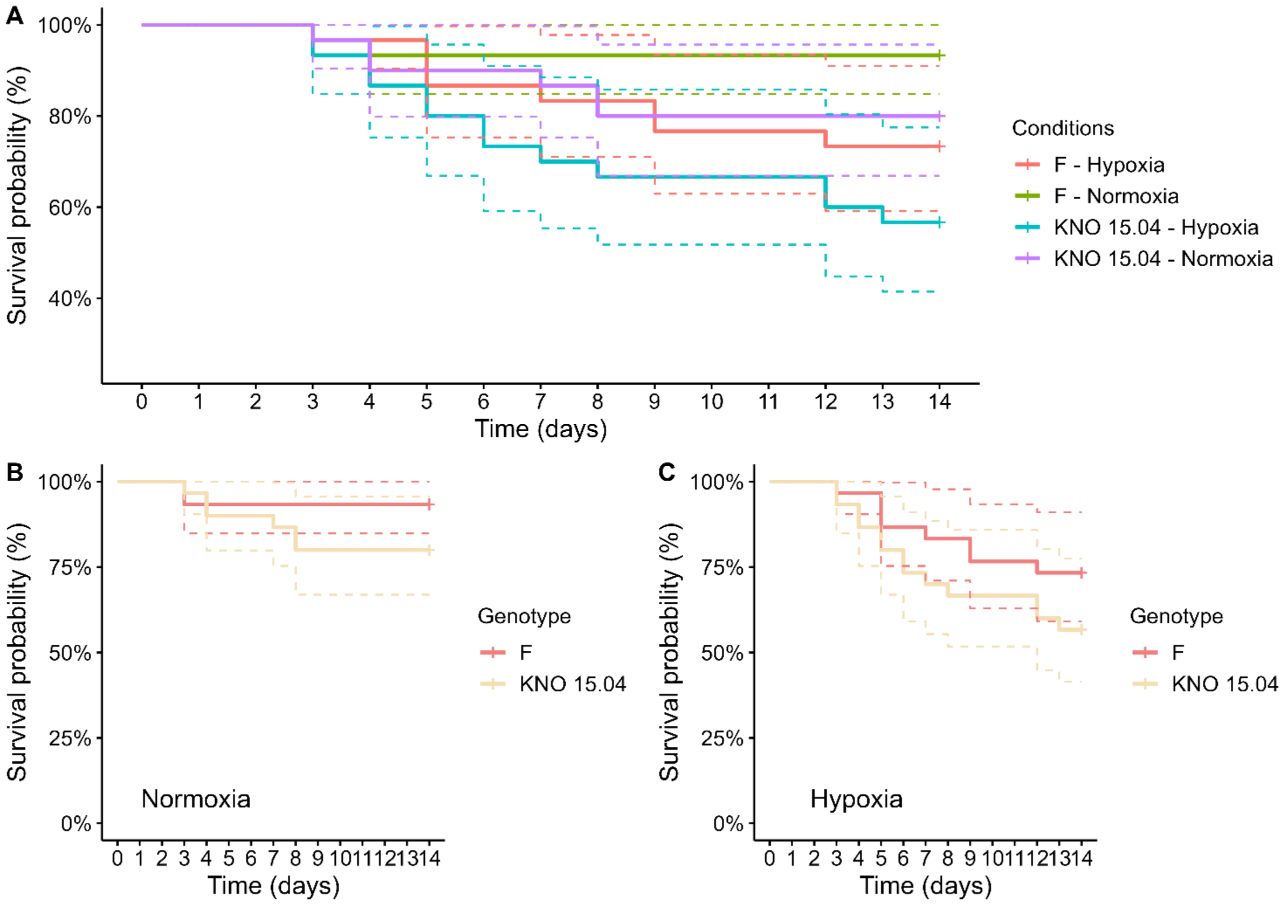


**Supplementary Figure 1.** Comparison of survival of KNO 15.04 and F genotypes in a hypoxia or normoxia environment. A) Full comparison of the two genotypes and two exposures. B-C) Comparison of the F and KNO 15.04 genotypes in normoxia and hypoxia, respectively. Dashed lines represent the 95% confidence intervals. Sample size was n=30 (10 individuals * 3 replicates) for each genotype * exposure combination.

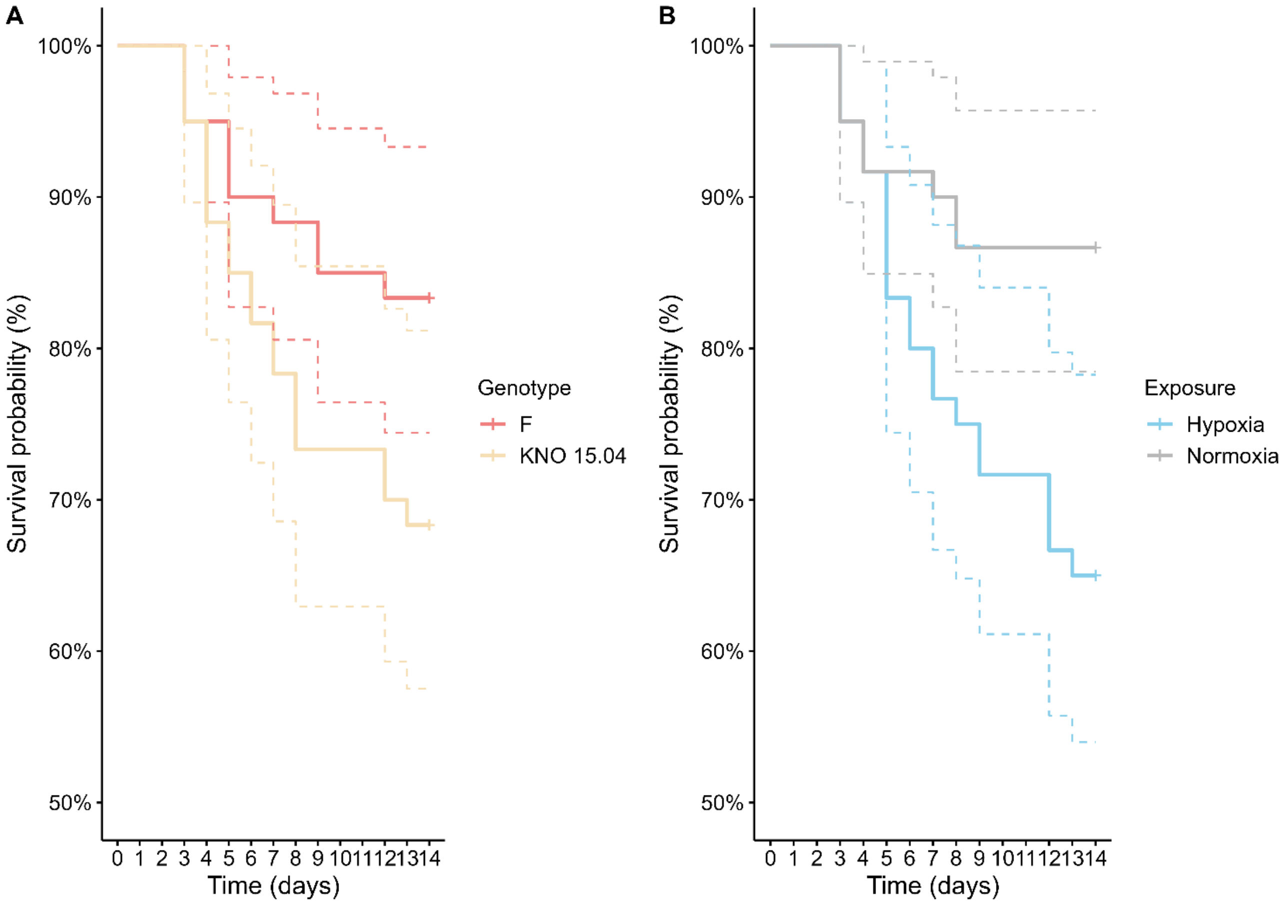


**Supplementary Figure 2.** Comparison of survival of KNO 15.04 and F genotypes in a hypoxia or normoxia environment. A) Comparison of F and KNO 15.04 genotypes in both exposures. Red lines correspond to survival of the F genotype and orange lines to survival of the KNO 15.04 genotype. B) Comparison of hypoxia and normoxia exposure over both genotypes. Grey lines correspond to survival in a normoxia exposure and blue lines to survival in a hypoxia exposure. Dashed lines represent the 95% confidence intervals. Sample size was n=30 (10 individuals * 3 replicates) for each genotype * exposure combination.

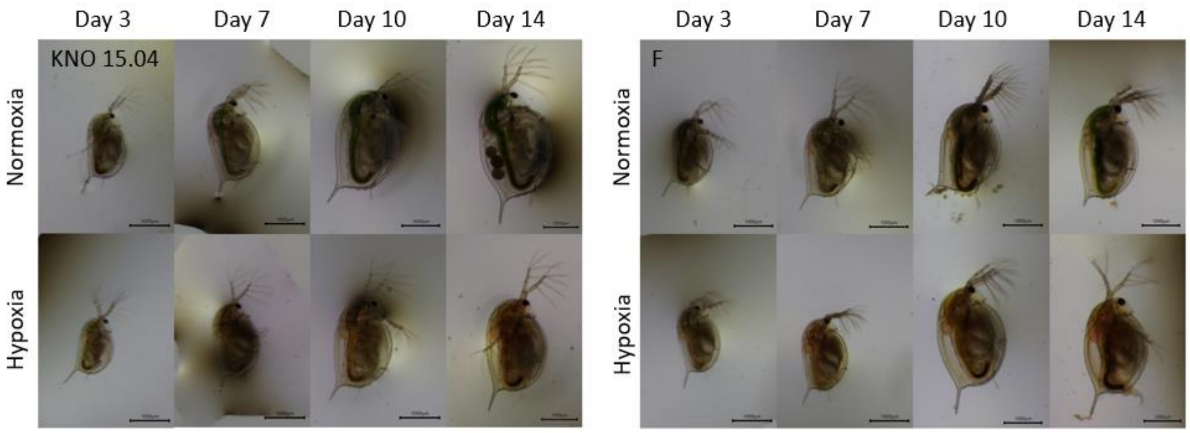


**Supplementary Figure 3.** Phenotypic appearance over time of normoxia (upper panels) and hypoxia (bottom panels) exposed D. magna of the KNO 15.04 (left figures) and F (right figures) genotype.

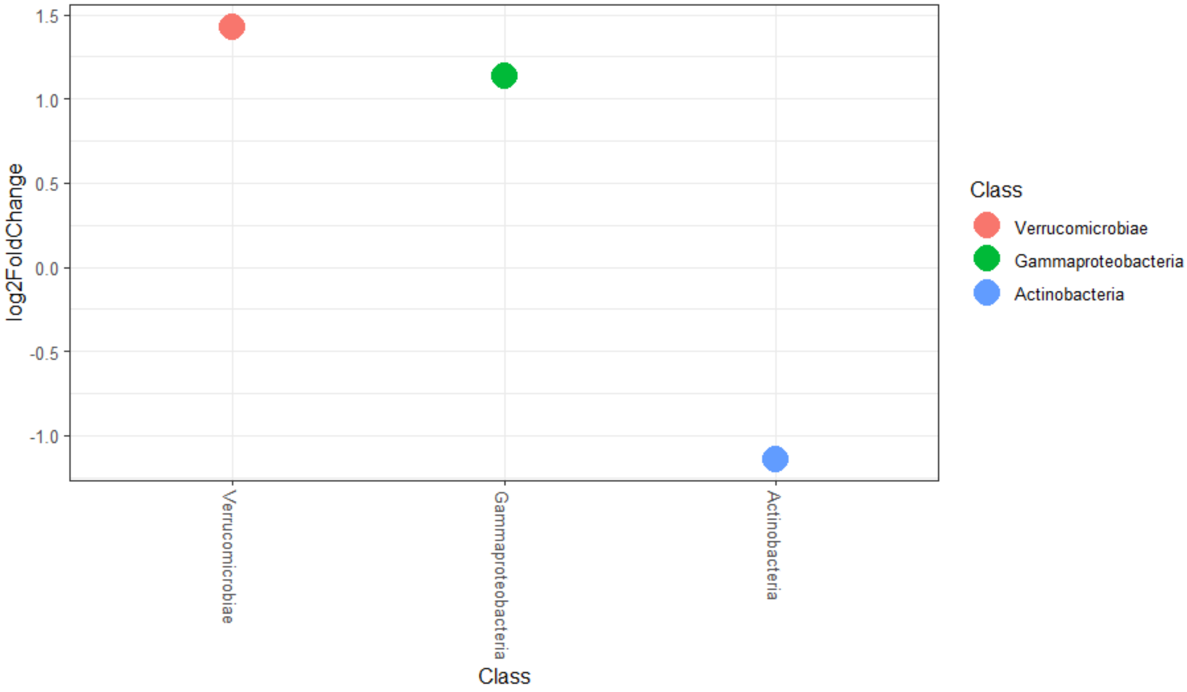


**Supplementary Figure 4.** Deseq2 comparison of classes between normoxia and hypoxia in the bacterioplankton samples with negative log2FoldChange values of bacterial classes being less represented in normoxia.

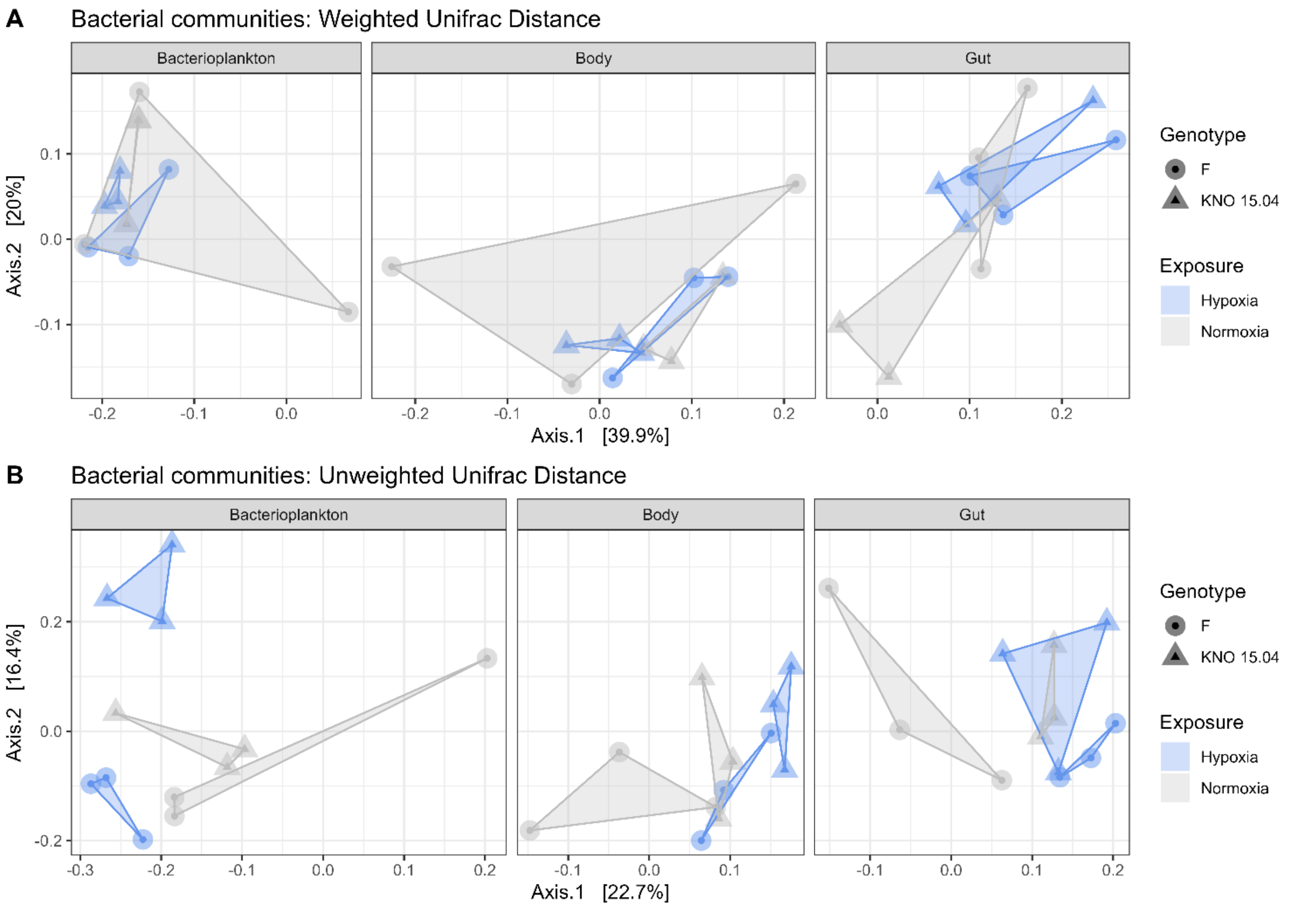


**Supplementary Figure 5**. Beta diversity: (A) Weighted and (B) Unweighted Unifrac distance of the bacterial communities within the different sample types (left panel: bacterioplankton, middle panel: body, right panel: gut). Circles correspond to the F genotype, triangles to the KNO 15.04 genotype. Hypoxia exposure is depicted in blue, normoxia exposure in grey.

**Supplementary Table 1.** Relative mean class counts per sample type x exposure combination. Grey marked and bold numbers are significant different numbers, bold numbers without marking are not significant but interesting numbers.

| | **Bacterioplankton** | | **Body** | | **Gut** | |
|---|---|---|---|---|---|---|
| **Class** | **Hypoxia** | **Normoxia** | **Hypoxia** | **Normoxia** | **Hypoxia** | **Normoxia** |
| Bacteroidia | 49,007452 | **48,15794703** | 58,47184908 | 55,96729321 | 27,54841169 | **44,77306509** |
| Gammaproteobacteria | **12,80593703** | **16,24254468** | 35,56615448 | 25,7845279 | 48,67385049 | 38,42611634 |
| Actinobacteria | **24,59862856** | **10,08552273** | 0,654258523 | 3,120085215 | 6,699315313 | 3,28761547 |
| Verrucomicrobiae | **10,91930858** | **23,46311698** | 0,456096548 | **2,045449712** | 2,824877433 | **2,367270962** |
| Alphaproteobacteria | 1,650147826 | 1,573801194 | 3,274537082 | 12,09568286 | 10,74472269 | 10,64480521 |
| Bdellovibrionia | 0,537753415 | 0,131309996 | 1,50338234 | 0,457449245 | 3,389427019 | 0,235581352 |
| Polyangia | 0,038998456 | 0,098920028 | **0** | 0,336156013 | **0** | 0,102064544 |
| Gracilibacteria | 0,076225725 | 0,031985075 | 0,073721944 | 0,186336822 | 0,073772256 | 0,036903402 |
| Planctomycetes | 0,141413974 | 0,170917705 | 0 | 0,007019025 | 0,007019211 | 0,063235743 |
| Bacilli | 0,090206328 | 0 | **0** | **0** | 0,026322444 | 0,005266484 |
| Desulfitobacteriia | 0,065214918 | 0 | **0** | **0** | 0,012281441 | 0,00351395 |
| Acidimicrobiia | 0,05989578 | 0,008780248 | **0** | **0** | 0 | 0,001753463 |
| Armatimonadia | 0 | 0 | **0** | **0** | 0 | 0,049294039 |
| Kapabacteria | 0 | 0,035154327 | **0** | **0** | 0 | 0,00351395 |
| Oligoflexia | 0,008817409 | 0 | **0** | **0** | 0 | 0 |
